# ProtSEC: Ultrafast Protein Sequence Embedding in Complex Space Using Fast Fourier Transform

**DOI:** 10.1101/2025.08.17.670693

**Authors:** Rajan Saha Raju, Rashedul Islam

**Affiliations:** Department of Computer Science and Engineering, Shahjalal University of Science and Technology, Sylhet-3114, Bangladesh; Omics-Lab, Bioinformatics and Computational Biology; Environmental Genomics & Systems Biology Division, Lawrence Berkeley National Laboratory, Berkeley, CA, USA

**Keywords:** embedding, BLOSUM62, complex number, fast fourier transform, protein sequence

## Abstract

Among the various approaches for representing protein sequences as vectors, embeddings derived from protein language models (PLMs) have been empirically shown to enhance accuracy in downstream bioinformatics tasks. However, the substantial computational demands of PLMs, both during training and inference, pose significant challenges. We introduce ProtSEC (**Prot**ein **S**equence **E**mbedding in **C**omplex Space), a novel approach that encodes each amino acid as a unique complex number derived from the BLOSUM62 substitution matrix. By modeling protein sequences as complex signals and applying the Fast Fourier Transform (FFT), ProtSEC generates embeddings in the complex space. Unlike PLMs, ProtSEC requires no pre-training on large protein sequence datasets and operates independently of any pre-trained models. Our benchmarking demonstrate that ProtSEC achieves a 20,000-fold reduction in runtime and an 85-fold improvement in memory efficiency compared to popular PLMs (e.g., esm2_3B, esm2_35M, prot_t5, prot_bert). Depending on the task, ProtSEC demonstrates either superior or comparable accuracy to PLMs in sequence similarity search, sequence classification and phylogenetic tree reconstruction. ProtSEC provides fast and accurate protein sequence embeddings in complex numbers, facilitating efficient integration into diverse downstream bioinformatics workflows. ProtSEC is available at https://github.com/omics-lab/ProtSEC.

## Introduction

Protein sequence embedding refers to the transformation of protein sequences into numerical vector representations within a continuous space. These embeddings are typically high-dimensional (e.g., 512, 1,024, or even 2,048 dimensions), where similar proteins are mapped close together in the vector space. Among different techniques for vector representation of protein sequences, amino acid letters are often directly encoded into discrete numerical representations using their alphabetic, physicochemical and evolutionary features^1–4^. Discrete encoding methods may not capture complex interactions among amino acids, structural patterns, and positional dependencies. In contrast, embedding methods generate continuous vector representations that capture functional and structural properties of protein sequences. Currently, a large number of learning-based protein language models (PLMs) are available that could effectively learn sequence syntax, folding and contextual information depending on the model; examples include SeqVec^5^, ProtVec^6^, TAPE^7^, UniRep^8^, CARP^9^, ProteinBERT^10^, ESM2^11^, and ProtTrans^12^. There are several variants of those PLMs which vary in the size of training datasets, number of parameters, accuracy, computational cost, and application needs. The accuracy of PLM generally depends on the size of the models, where larger models tend to have higher accuracy^11^. The key challenges in PLMs include: 1) the need for large and comprehensive training datasets, as different PLMs are often trained on diverse protein sets that may often represent biases and incompleteness; 2) the significant computational demands during both model training and the subsequent generation of embeddings for new sequences; and 3) limitations of embedding dimensionality, where representations in relatively low-dimensional spaces fail to capture the subtle distinctions required to separate the vast number of relevance groups^13^. These challenges significantly impede the integration of PLMs into downstream bioinformatics and machine learning workflows, especially when handling large protein sequence datasets. Thus, there’s a need for the development of protein sequence embedding methods that overcome those shortcomings of PLMs while offering reasonable accuracy in the downstream bioinformatics tasks.

Here we propose **ProtSEC** (**Pro**tein **S**equence **E**mbedding in **C**omplex Space), which maps amino acids to complex numbers based on evolutionary relationships captured in the BLOSUM62^4^ and transforms complex signals of protein sequences into fixed-length complex-valued embedding vectors using the Fast Fourier Transform (FFT)^14^, a faster version of the Discrete Fourier Transform (DFT) algorithm. This approach in ProtSEC addresses major limitations of current PLMs by offering 1) independence from pre-training on large protein datasets, 2) computational efficiency and speed, and 3) flexibility to choose the embedding dimension based on the distribution of protein sequence lengths. DFT and FFT have previously been applied to the phylogenetic analysis of protein sequences^15,16^ and identifying functional regions^17^ using physicochemical properties of AAs. The FFT was used in MAFFT^18^ for multiple sequence alignment to rapidly identify homologous regions by efficiently calculating the correlation between two sequences. While previous methods showed the potential of DFT in extracting evolutionary and functional properties as frequency components from protein sequence, ProtSEC has been tested with diverse datasets for sequence similarity search, sequence classification and phylogenetic tree reconstruction. Benchmarking against popular PLMs revealed ProtSEC achieves comparable accuracy to popular PLMs while overcoming key challenges of PLMs. ProtSEC enables embedding of large protein sequence datasets on a personal computer, representing a significant advancement in protein sequence embedding by facilitating faster data processing and accelerating discoveries in protein language understanding.

### Results

### Overview of ProtSEC protein sequence embedding architecture

We used BLOSUM62 substitution matrix^4^ as the basis to project the AAs into a complex geometric space using Multi Dimensional Scaling (MDS) **(Figure 1A)**. Complex-valued representations of amino acids carried substantially more informative features than real-valued ones. Compared to one-dimensional real-values, two-dimensional complex-values achieved significantly higher precision in preserving evolutionary relationships in low-dimensional projection with 65.7% reduction of stress (from 12,424 to 4,263) in MDS estimated by Kruskal’s stress^19^. Additionally, projection of AAs into the complex space revealed meaningful patterns that correspond to their underlying biochemical properties. Hydrophobic amino acids formed a distinct cluster in the complex plane, while ambiguous codes such as B and Z were positioned in the middle of their respective related AAs, B between Aspartic acid (D) and Asparagine (N) and Z between Glutamic acid (E) and Glutamine (Q). Cysteine (C) and Tryptophan (W) appeared more distant from the other AAs, consistent with their unique characteristics: Cysteine’s ability to form disulfide (S–S) bonds and Tryptophan’s highest molecular weight among the standard AAs. These observations demonstrate that the complex number representation effectively captures distinct biochemical properties of amino acids. Thus, we encoded each protein sequence as a complex signal using the precomputed complex representations of the 22 amino acids. The FFT was then applied to convert the complex signal of arbitrary length from the spatial domain (linear arrangement of amino acid residues) into the spectral domain **(Figure 1B, Methods)**. This transformation captures periodic and structured sequence patterns, generating a fixed-length complex-valued embedding vector. We evaluated ProtSEC’s accuracy using various dimensionality reduction methods (MDS, t-SNE, UMAP) and distance functions (SMS, ASMP, SNN) with 1024-dimensional embeddings. The MDS-ASMP combination yielded the highest accuracy in sequence similarity search **(Figure S1)**. In ProtSEC, the embedding dimension is adjustable, allowing for examination of the sequence length distribution to determine optimal dimensionality. In contrast, PLM-based models offer no flexibility to modify embedding dimensions after training. Representations constrained to relatively low-dimensional spaces are insufficient for capturing the subtle distinctions required to separate the vast number of relevance groups^13^.

**Figure 1:**
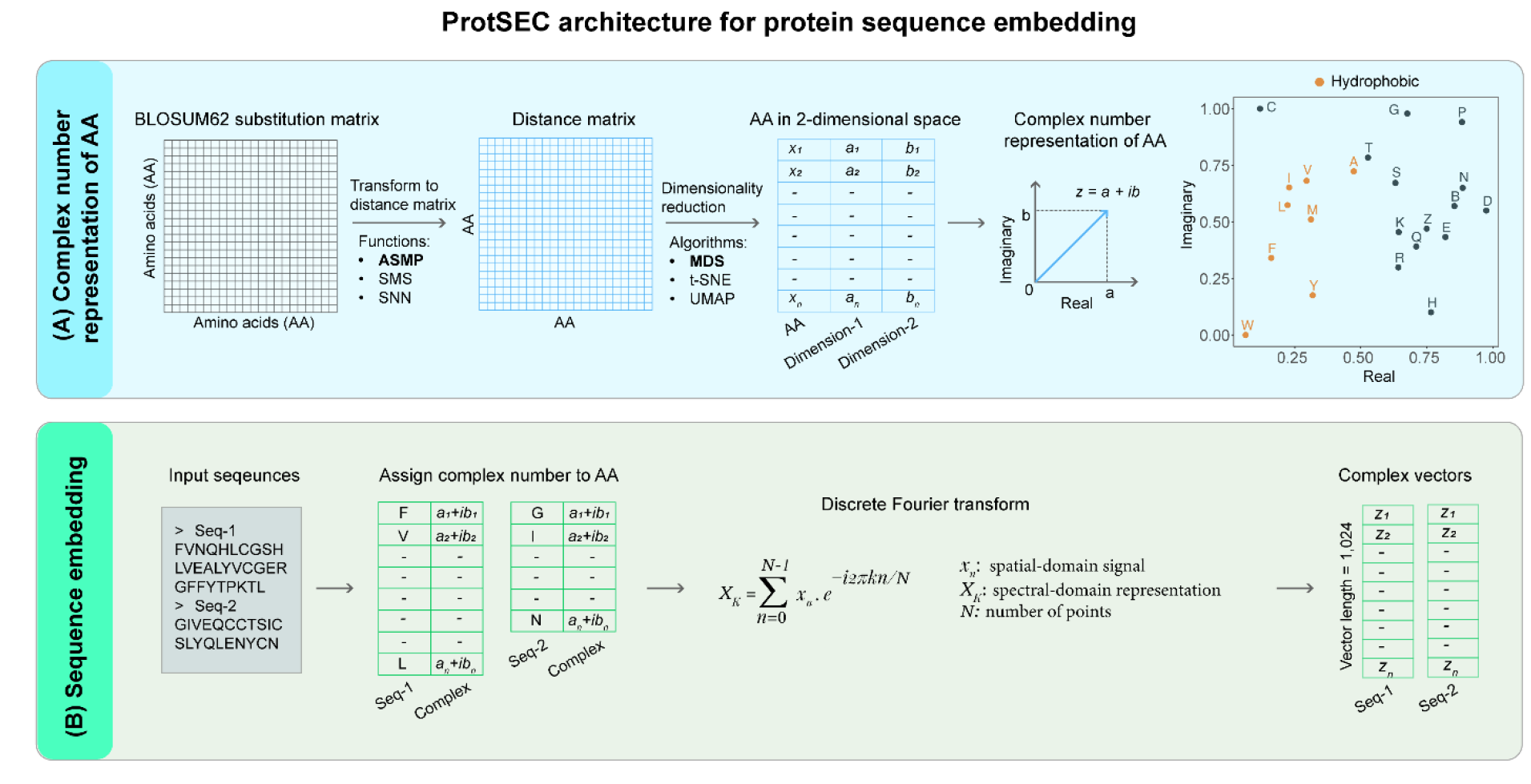
Overview of ProtSEC architecture. **A)** ProtSEC architecture for complex number representation of amino acids (AA). BLOSUM62 substitution matrix is transformed to a distance matrix using one of the three distance functions such as Average of Self Minus Pairwise (ASMP), Subtraction from Max Similarity (SMS), and Shift and Normalize (SNN). Then amino acids are projected into a two-dimensional space from their distance matrix using either Multi-Dimensional Scaling (MDS), t-distributed Stochastic Neighbor Embedding (t-SNE), or Uniform Manifold Approximation and Projection (UMAP). In the scatter plot, the first component is represented as the real part and the second component as the imaginary part of a complex number for each amino acid (n=22, including ambiguous codes). **B)** Each amino acid residue in the protein sequence is mapped to its corresponding complex number which was precomputed in the previous step. The complex sequence is then passed through Discrete Fourier Transform (DFT) to generate a complex-valued embedding vector.

### ProtSEC outperformed PLMs in sequence similarity search

To benchmark the accuracy of ProtSEC in finding biological similarity between sequences, we compared ProtSEC with the widely recognized reference method BLASTp (Basic Local Alignment Search Tool for proteins)^20^ and PLM-based embedding methods. BLASTp aligns a query sequence to the sequence database and then scores these alignments using a substitution scoring matrix (such as BLOSUM or PAM matrices)^20^. For PLM-based embedding methods, we calculated the cosine similarity between the query sequence vector and the precomputed database of sequence vectors to identify the most similar matches **(Figure S2)**. We used two popular smaller PLMs (prot_bert with 420 millions of parameters and esm2_35M with 35 millions of parameters) and two large PLMs (esm2_3B and prot_t5 both with 3 billions of parameters)^11,12^. To evaluate the accuracy of sequence similarity search of different embedding methods, we queried the 5K dataset against 5K, 10K, 20K, and 40K protein sequences datasets **(Table S1)**. Those benchmark datasets contain well characterized protein sequences of UniProtKB/Swiss-Prot with average length of ∼370 amino acid residues^21^. As expected, querying the 5K dataset against itself resulted in correct identification (100% accuracy) across all methods. Querying the 5K dataset against rest of the datasets showed BLASTp consistently outperformed other methods, achieving an average accuracy of 71.83%, where prot_bert performed worst with average accuracy of 60.24%. The accuracy of all methods improved with increasing dataset size, attributable to the higher likelihood of retrieving similar sequences from larger datasets **(Table 1)**. In sequence similarity search, ProtSEC is 2.44% less accurate compared to BLASTp but demonstrated 0.49% and 9.15% higher accuracy compared to the two smaller PLMs such as esm2_35M and prot_bert, respectively. While the two large-scale PLMs exhibited an average of 0.33% higher accuracy than ProtSEC (esm2_3B showed a 0.26% increase and prot_t5 achieved a 0.39% increase), the improvements were marginal.

**Table 1:** Performance comparison for protein sequence similarity search. Accuracy is shown in percentages.

| Dataset | BLASTp | ProtSEC | esm2_3B | esm2_35M | prot_t5 | prot_bert |
| --- | --- | --- | --- | --- | --- | --- |
| 5K | 100 | 100 | 100 | 100 | 100 | 100 |
| 10K | 56.24 | 52.48 | 54.08 | 52.24 | 52.94 | 35.4 |
| 20K | 63.06 | 59.50 | 60.76 | 59.04 | 60.32 | 47.12 |
| 40K | 68.02 | 65.58 | 63.74 | 64.34 | 65.86 | 58.42 |
| **Average** | **71.83** | **69.39** | **69.65** | **68.91** | **69.78** | **60.24** |

### Evaluation of ProtSEC in sequence classification and phylogenetic tree reconstruction

We benchmarked ProtSEC for classifying proteins by their Enzyme Commission class 1 (EC1) numbers using the CARE dataset^22^. We used two test datasets consisting of 560 (30-50% identity to the training sequences) and 432 (<30% identity to the training sequences) sequences. We have generated two training datasets consisting of 5,000 and 2,000 sequences by random sampling of sequences from 184,529 CARE training dataset. While models trained with the entire CARE training dataset^22^ (n = 184,529 sequences) showed higher accuracy, embedding of large datasets using PLMs requires significant computation and run time. Therefore, we focused on assessing the relative accuracy using smaller training datasets of 5,000 and 2,000 sequences across embedding methods. We generated embeddings for both query and training datasets using ProtSEC and four PLMs, which were utilized as feature vectors for the classifier model. To evaluate performance, we employed K-Nearest Neighbors (KNN) classification^23^ with K=5 using five-fold cross-validation and reported the average accuracy across all folds. As anticipated, all methods exhibited superior EC1 classification accuracy for the query dataset with 30-50% identity to the training dataset compared to that with <30% identity, consistently across two training sets. The accuracy trends were similar between two training sets suggesting that the sampling of these sets did not introduce bias into the relative accuracy measurements. ProtSEC demonstrated an average accuracy that was 4.61% higher than esm2_3B, esm2_35M, and prot_bert when two query datasets were classified using KNN (**Figure 2A)**. However, the average accuracy of ProtSEC was 5.85% lower than prot_t5. These findings suggest that KNN-based classification using ProtSEC embedding demonstrates superior overall accuracy compared to PLMs. Nevertheless, it is imperative to acknowledge that the optimal embedding methodology may differ based on the specific requirements of the task.

**Figure 2:**
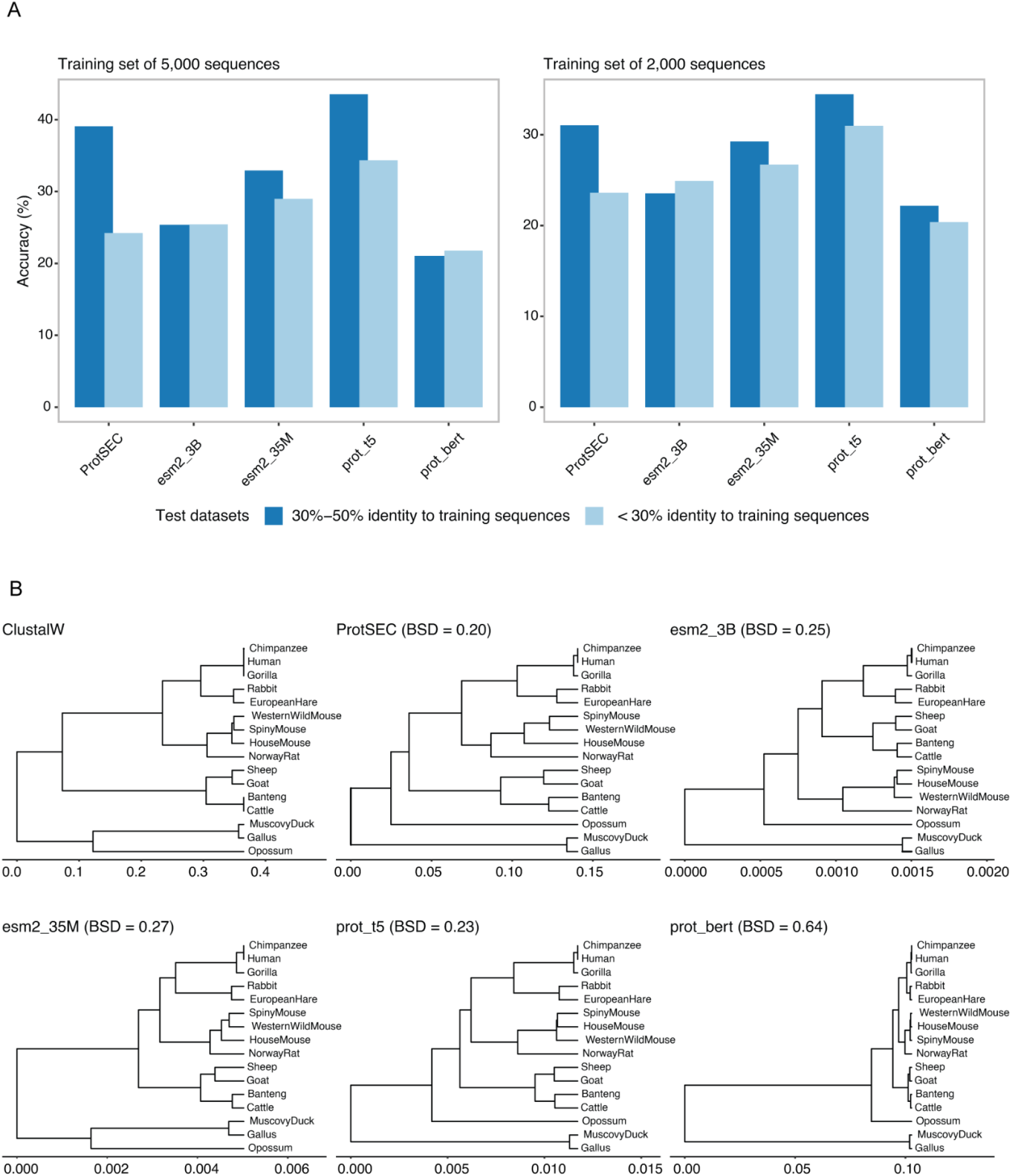
Phylogenetic reconstruction and clustering performance of ProtSEC. **B)** Accuracy in classifying Enzyme Commission class 1 for embedding methods. Two sets of query sequences have different sequence identities to the reference databases. **B)** Phylogenetic trees of the 16-BetaSet dataset reconstructed using ClustalW (multiple sequence alignment), ProtSEC, and PLM-based embeddings (esm2_3B, esm2_35M, prot_t5, prot_bert). Values on the X-axis of a tree represent branch lengths from the root. For BSD, trees were normalized by scaling all branch lengths proportionally.

We further evaluated ProtSEC on phylogenetic tree reconstruction using beta-globin sequences from different species; the 10-BetaSet (10 species) and 16-BetaSet (16 species) datasets^15^. We compared the performance of ProtSEC against a multiple sequence alignment–based reconstruction with ClustalW and PLM embeddings, including esm2_35M, esm2_3B, prot_t5 and prot_bert (**Figures 2B**,**S3)**. Using the ClustalW-based tree as reference, we calculated the normalized Branch Score Distance (BSD) for each method^24^. ProtSEC consistently achieved the lowest BSD compared to all PLM-based embeddings with two-fold less BSD compared to PLMs on average, indicating that its reconstructed trees were more topologically similar to the ClustalW reference. A notable difference emerged in the placement of Opossum. Consistent with previous study^15^, ClustalW clustered Opossum closer to Phasianoidea superfamily members (e.g., Gallus and Muscovy duck), whereas all PLM-based embeddings, except esm2_35M, grouped Opossum with mammals, which is taxonomically correct. esm2_35M reproduced the misplacement seen with ClustalW. Across all methods, the Bovoidea superfamily (e.g., goats, sheep, banteng, and cattle) consistently formed a distinct cluster away from Muroidea (spiny mice, western wild mice, house mice, and Norway rats), Hominidae (e.g., humans, chimpanzees, and gorillas), and Leporoidea (e.g., rabbits and European hares), except for the esm2_3B model. These suggest that despite some variations, embedding-based methods generally captured known taxonomies in these two datasets, where ProtSEC embedding provided more accurate tree reconstruction and achieved lower BSD values than the PLMs. ProtSEC’s robustness in aligning with known biology is likely due to its foundation being based on the evolutionary matrix, which appears to result in more reliable evolutionary separation.

### Computing performance evaluation of ProtSEC

Embedding of protein sequences using PLMs is a compute-intensive operation, and the requirement for computing resources depends on model size^11^. We benchmarked the computing performance by embedding the 5K dataset using ProtSEC, esm2_35M, esm2_3B, prot_bert and prot_t5 **(Methods, Table 2)**. ProtSEC does not depend on large pretrained models and relies on the precomputed file (9.7KB in size) containing complex number representation of amino acids. This precomputed file of ProtSEC is ∼14,000X smaller than the binary model file of esm2_35M, which is the smallest model we used in this benchmarking. Other models such as esm2_3B, prot_bert, and prot_t5 use model files several times larger than esm2_35M **(Table 2)**. Dependencies on a smaller precomputed file of ProtSEC accelerate the sequence embedding process markedly by both with reduced RAM and time requirements. ProtSEC showed ∼85X lower peak RAM usage and 20,000X lower computing time compared to PLMs on average. The peak RAM usage and the time generally increase with the increase of model sizes in PLMs.

**Table 2:**
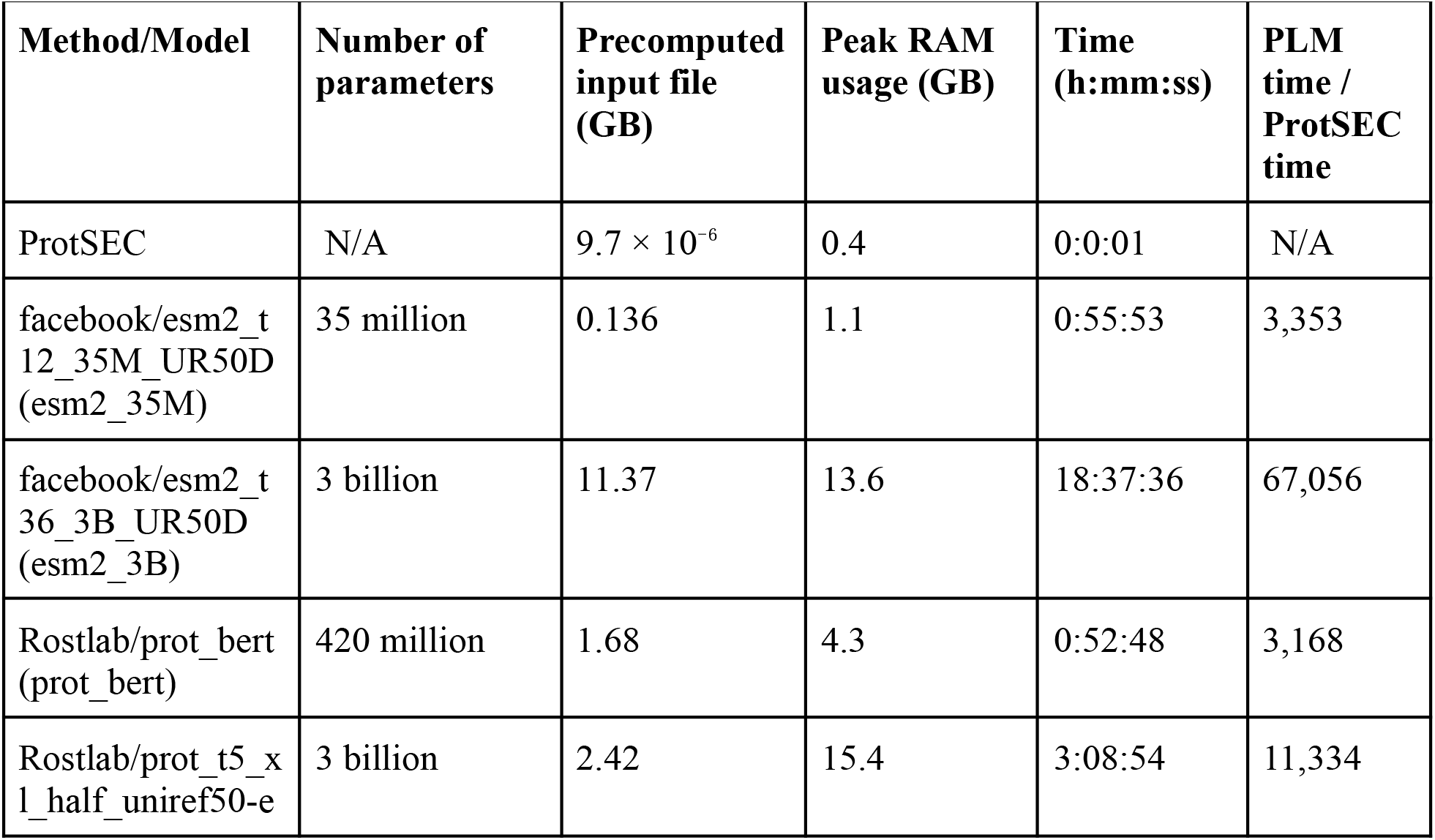
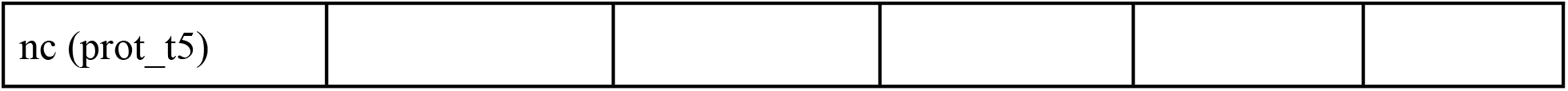
Benchmarking of computing performance of different protein sequence embeddings.

## Discussion

ProtSEC leverages complex-valued representation of amino acids that accurately captured evolutionary relationships and enabled DFT to extract richer spectral-domain features for characterizing both periodic and global sequence patterns. ProtSEC provides a comparable performance to the large pre-trained PLMs, without requiring extensive computing resources or run time. ProtSEC was able to recognize meaningful similarities between proteins, performing almost as well as traditional alignment methods and often better than some widely used PLMs. When tested on larger protein datasets, ProtSEC shows increasing accuracy in finding related sequences. In building evolutionary trees or classifying sequences based on EC1 class, ProtSEC showed a comparable accuracy to the established reference methods and PLMs, suggesting it can more reliably be used in the downstream bioinformatics applications. At the same time, ProtSEC was ∼20,000x faster and 85x leaner, showing how defining variables and method selection can greatly simplify a resource-intensive task. Another key advantage of ProtSEC is that it offers adjustable embedding dimensions, unlike PLM-based models. This flexibility allows for optimizing dimensionality based on sequence length and specific tasks, thereby addressing the limitation of low-dimensional representations, which often fail to capture subtle distinctions required to distinguish among relevance groups. Overall, our benchmarking suggests that ProtSEC offers a strong balance between biological insight and computational efficiency, making it a useful alternative to PLM-based embedding.

Our analysis suggests that no single model consistently outperformed others across all tested bioinformatics applications; performance varied depending on the specific task. Esm2_3B and esm2_35M showed variable accuracy depending on the database size such as esm2_35M outperformed esm2_3B for 40K dataset. Esm2_3B consistently outperformed esm2_35M in phylogenetic tree reconstruction. However, despite being a significantly larger model, esm2_3B showed lower accuracy in EC1 classification compared to esm2_35M. The prot_t5 consistently showed superior performance in sequence similarity and EC1 classification but not in the phylogenetic tree reconstruction. Despite those variations depending on the embedding methods, we have demonstrated ProtSEC embedding’s utility in several bioinformatics applications, further research is needed to implement and benchmark ProtSEC in a broader range of bioinformatics contexts. Since ProtSEC is based on the static evolutionary matrix, BLOSUM62, it may not be aware of context specific differences. Context aware embedding methods based on LPMs could be more suitable for context specific inferences. In our future research, PortSEC could be improved by providing context specific matrices instead of BLOSUM62.

Unlike PLM-based methods that produce real-valued vectors, ProtSEC generates complex-valued vectors, making direct classification benchmarking difficult. Most of the existing vector databases for real number support metrics such as Euclidean distance, cosine similarity and inner product^25^. However, none of the current solutions natively support complex-valued vectors. This limitation highlights the need for extending vector database capabilities to complex domains, where metrics such as the Hermitian inner product and phase correlation are essential. Addressing this gap would open new research opportunities and facilitate applications that rely on complex vector representations.

Implementing PLMs requires significant computational resources and often need access to GPU, and requires trained individuals to work with those models. These could impede the research outcome, especially considering that the users are exceptionally multidisciplinary: different domains might have different levels of computational skills when it comes to the implementation and integration of PLMs. In addition, smaller models tend to be downloaded more frequently compared to the larger models, despite their potential lower accuracy. This suggests the smaller models are popular due to their ease in use such as less requirements of computing resources and run time. Its self-contained nature and straightforward implementation made ProtSEC an attractive option for researchers for embedding large protein sequence datasets without the overhead or complex setups associated with PLMs or conventional methods.

## Methods

### Dataset

We have downloaded complete UniProtKB/Swiss-Prot protein sequences (n=572,970) on February 5th, 2025 from (https://ftp.expasy.org/databases/uniprot/current_release/knowledgebase/complete/uniprot_sprot.fasta.gz). From these, we selected 506,185 well-characterized proteins and then we randomly chose subsets of 5,000, 10,000, 20,000 and 40,000 sequences to create 5K, 10K, 20K, and 40K datasets (**Table S1**). CARE^22^ dataset was downloaded from (https://github.com/jsunn-y/CARE). Two groups of test sequences were used: 432 held-out sequences with <30% identity to the training set and 560 held-out sequences with 30–50% identity.

### Protein language model (PLM)

We used four widely used pre-trained PLMs such as esm2_3B, esm2_35M, prot_t5, prot_bert (**Table 1**) for embedding^11,12^. All models are publicly available at Hugging Face (**Table S2**).

### ProtSEC embedding algorithm

#### 1. Computation of a pairwise distance matrix D

The BLOSUM62 matrix tells the substitution score for replacing one amino acid with another. Formally, this matrix, denoted as *S*, contains values *S*[*i*][*j*] indicating the substitution similarity between amino acids *i* and *j*. To transform these similarity scores into a distance measure, we compute a pairwise distance matrix *D* using one of the three distance functions mentioned below, each designed to reinterpret similarity into distance in a distinct way:

##### 1.1 Average of Self Minus Pairwise (ASMP)

The ASMP function defines the distance between amino acids *i* and *j* as:

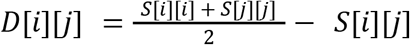

This function interprets distance as the difference between the average self-similarity of the two amino acids and their direct pairwise similarity. An average larger self-similarity relative to their mutual similarity increases the distance.

##### 1.2. Subtraction from Max Similarity (SMS)

The SMS function computes distance by subtracting the pairwise similarity from the maximum similarity observed in the entire matrix. This approach scales distances relative to this maximum.

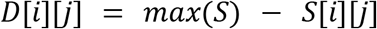

This approach treats the most similar amino acid pair as having zero distance and scales all other distances relative to this maximum. It effectively inverts the similarity scale to define distances.

##### 1.3. Shift and Normalize (SNN)

The SNN method converts a similarity matrix into a distance matrix where distances are scaled between 0 and 1. This involves two key steps: shifting and normalizing.

###### 1.3.1 Shifting to Non-negative Values

To remove negative values of the BLOSUM62 matrix, we shift the matrix by subtracting the minimum value of the matrix from every element. After shifting, the smallest entry in S′ becomes 0, and all other entries are non-negative.

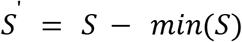

###### 1.3.2 Normalizing to [0, 1] Range

The shifted similarities are normalized so that they fall within the range [0, 1]. This is achieved by dividing each entry by the maximum value in the shifted matrix:

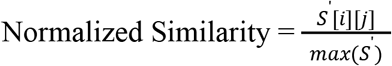

###### 1.3.3 Converting Similarity to Distance

To transform similarities into distances (where higher similarity means lower distance), we invert the normalized similarities:

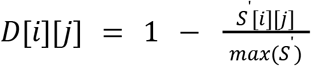

##### 2. Dimensionality reduction for complex number encoding

Given the computed distance matrix D, we employ dimensionality reduction techniques such as Multidimensional Scaling (MDS)^26^, t-Distributed Stochastic Neighbor Embedding (t-SNE)^27^, or Uniform Manifold Approximation and Projection (UMAP)^28^ to project each amino acid into a two-dimensional space. Each amino acid (n = 22, including standard and ambiguous codes) is then represented as a complex number, where the first dimension of the projection corresponds to the real part and the second dimension corresponds to the imaginary part. This mapping establishes a unique complex representation for each amino acid.

##### 3. Embedding by Discrete Fourier Transform

For a given protein sequence, each residue is encoded using its corresponding complex number, resulting in a vector of complex numbers. To obtain the embedding, we apply the Discrete Fourier Transform (DFT) to the vector of complex numbers, projecting it into a complex frequency domain. The resulting transformed sequence serves as the embedding of the protein in a complex-valued feature space. Mathematically, the DFT of a sequence *x* = [*x*_0_, *x*_1_, …, *x*_*N*−1_] is defined as:

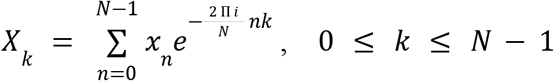

where N is the length of the sequence, *x*_*n*_ denotes the n-th complex-valued residue encoding, and *X*_*k*_ represents the transformed value in the frequency domain. The resulting transformed sequence *X* = [*X*_0_, *X*_1_, …, *X*_*N*−1_] serves as the embedding of the protein in a complex-valued feature space. To efficiently compute the DFT, we utilize the Fast Fourier Transform (FFT)^14^, which significantly reduces the computational complexity from *O*(*N*^2^) to *O*(*N log*(*N*)), enabling scalable processing of protein sequences.

### Sequence similarity search

Using protein language models (PLMs), we generate embedding vectors for all protein sequences in the dataset to construct a vector database. For a given query sequence, its embedding vector is generated and computed cosine similarity against the precomputed vectors in the vector database. The sequence with the highest similarity score is then retrieved as the closest match. For two real vectors *x,y* ϵ ℝ^*n*^, the cosine similarity is:

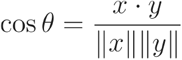

For normalized complex vectors, the modulus of the Hermitian inner product extends the concept of cosine similarity from real to complex domains. In ProtSEC, we adopt phase correlation as the similarity metric, rather than relying on the Hermitian inner product, as it demonstrates superior performance. For two FFT-transformed complex vectors *x,y* ϵ ℂ^*n*^, the phase correlation score is:

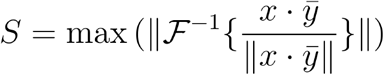

Here, ℱ^-1^ and bar sign indicate inverse Fourier transform and complex conjugation respectively. For BLASTp (with default parameters), bit scores were used to rank hits. Based on bit scores for BLASTp and similarity scores for embedding methods, the highest-ranked hit was selected in each case, and accuracy was evaluated by matching the Gene Name (GN) identifiers between query and retrieved sequences. The implementation of the sequence similarity search pipeline using the four PLMs is available at https://github.com/Rajan-sust/GeneAnnotation.

### Phylogenetic tree reconstruction

Phylogenetic trees were constructed using UPGMA clustering^29^ on distance matrices derived from multiple protein representation methods. For sequence-based analysis, protein sequences were aligned using ClustalW, and evolutionary distances were calculated using maximum likelihood estimation. For protein language model (PLM) approaches, pre-computed distance matrices were generated from protein embeddings using PLMs and ProtSEC methods. To enable quantitative comparison across methods, Branch Score Distance (BSD)^30^ was calculated between each embedding-derived tree and the ClustalW reference tree. BSD was performed using normalized tree topologies by scaling all branch lengths to sum to unity, ensuring that topological differences rather than absolute distance scales drove the comparisons.

### K-Nearest Neighbour Classification

For KNN classification, we utilized phase correlation distance (defined as 1.0 ™ phase correlation) for the ProtSEC embedding and cosine distance for the PLM-derived embedding as the distance metric. Classification was based on a majority vote, with random tie-breaking. In cases of ties for the Kth nearest vector, all tied candidates were included in the vote.

### Recommendations for Selecting the Embedding Dimension

The embedding vector length (d) determines the output length of the Fast Fourier Transform (FFT). Appropriate selection of d is essential for balancing computational efficiency with information preservation. The relationship between d and input sequence length follows three cases:

1. When d < input length, sequences are truncated from the end.
2. When d > input length, sequences are zero-padded at the end.
3. When d is unspecified, a default value of d = 1024 is applied.

For datasets exhibiting substantial variation in sequence lengths, we recommend setting d to the third quartile (Q3) of the length distribution. To optimize FFT computational performance, d can be rounded up to the nearest power of two (≥ Q3), as FFT algorithms achieve maximum efficiency at power-of-two lengths.

Implications:

1. Sequences exceeding d will be truncated, resulting in minor information loss.
2. Sequences shorter than d will be zero-padded.
3. This strategy represents a practical trade-off between computational speed and the marginal information loss incurred through truncation.

For datasets characterized by uniformly short sequences, d should be set to the maximum observed sequence length. This eliminates truncation entirely, ensuring complete information preservation. Rounding d up to the nearest power of two further enhances FFT efficiency.

Implications:

1. All sequences will be zero-padded to length d.
2. FFT computation remains efficient while preserving all original sequence information.

### Computing performance

To evaluate the computational performance of different embedding methods, we used 5K dataset in a desktop computer (CPU model: 12th Gen Intel(R) Core(TM) i9-12900KS, Total physical memory: 127.8GB, Storage device model: Samsung SSD 970 EVO Plus 2TB, Interface type: SCSI, Size: 2TB). We used elapsed (wall clock) time as embedding time and maximum resident set size as the peak RAM usage. We used the default parameters of embedding methods for the evaluation of computing performance.

## Data and code availability

Source code and dataset are available on GitHub at: https://github.com/omics-lab/ProtSEC

## Acknowledgments

This study did not receive any funding. We acknowledge Omics-Lab (https://omics-lab.com/) for providing the research platform.

## Author contributions

**Rajan Saha Raju** Conceptualization, Methodology, Software, Formal analysis, Investigation, Data curation, Visualization, Writing – original draft, Writing – review & editing.

**Rashedul Islam** Conceptualization, Methodology, Formal analysis, Investigation, Data curation, Writing – original draft, Writing – review & editing, Visualization, Supervision.

## References

1. Yang, K. K., Wu, Z., Bedbrook, C. N. & Arnold, F. H. Learned protein embeddings for machine learning. Bioinformatics 34, 2642–2648 (2018).

2. Harding-Larsen, D. et al. Protein representations: Encoding biological information for machine learning in biocatalysis. Biotechnol. Adv. 77, 108459 (2024).

3. Mei, H., Liao, Z. H., Zhou, Y. & Li, S. Z. A new set of amino acid descriptors and its application in peptide QSARs. Pept. Sci. 80, 775–786 (2005).

4. Henikoff, S. & Henikoff, J. G. Amino acid substitution matrices from protein blocks. Proc. Natl. Acad. Sci. 89, 10915–10919 (1992).

5. Heinzinger, M. et al. Modeling aspects of the language of life through transfer-learning protein sequences. BMC Bioinformatics 20, 723 (2019).

6. Asgari, E. & Mofrad, M. R. Continuous distributed representation of biological sequences for deep proteomics and genomics. PloS One 10, e0141287 (2015).

7. Rao, R. et al. Evaluating Protein Transfer Learning with TAPE. (2019).

8. Alley, E. C., Khimulya, G., Biswas, S., AlQuraishi, M. & Church, G. M. Unified rational protein engineering with sequence-based deep representation learning. Nat. Methods 16, 1315–1322 (2019).

9. Yang, K. K., Fusi, N. & Lu, A. X. Convolutions are competitive with transformers for protein sequence pretraining. Cell Syst. 15, 286–294.e2 (2024).

10. Brandes, N., Ofer, D., Peleg, Y., Rappoport, N. & Linial, M. ProteinBERT: a universal deep-learning model of protein sequence and function. Bioinformatics 38, 2102–2110 (2022).

11. Lin, Z. et al. Evolutionary-scale prediction of atomic-level protein structure with a language model. Science 379, 1123–1130 (2023).

12. Elnaggar, A. et al. ProtTrans: Toward Understanding the Language of Life Through Self-Supervised Learning. IEEE Trans. Pattern Anal. Mach. Intell. 44, 7112–7127 (2022).

13. Weller, O., Boratko, M., Naim, I. & Lee, J. On the Theoretical Limitations of Embedding-Based Retrieval. (2025).

14. Hufnagel, R. & Stanley, N. An algorithm for the machine calculation of complex Fourier series. J. Opt. Soc. Am. 54, 52–61 (1964).

15. Li, W. et al. FFP: joint Fast Fourier transform and fractal dimension in amino acid property-aware phylogenetic analysis. BMC Bioinformatics 23, 347 (2022).

16. Hou, W., Pan, Q., Peng, Q. & He, M. A new method to analyze protein sequence similarity using Dynamic Time Warping. Genomics 109, 123–130 (2017).

17. Ramachandran, P., Antoniou, A. & Vaidyanathan, P. P. Identification and location of hot spots in proteins using the short-time discrete Fourier transform. in Conference Record of the Thirty-Eighth Asilomar Conference on Signals, Systems and Computers, 2004. vol. 2 1656–1660 Vol.2 (2004).

18. Katoh, K., Misawa, K., Kuma, K. & Miyata, T. MAFFT: a novel method for rapid multiple sequence alignment based on fast Fourier transform. Nucleic Acids Res. 30, 3059–3066 (2002).

19. Kruskal, J. B. Multidimensional scaling by optimizing goodness of fit to a nonmetric hypothesis. Psychometrika 29, 1–27 (1964).

20. Altschul, S. F., Gish, W., Miller, W., Myers, E. W. & Lipman, D. J. Basic local alignment search tool. J. Mol. Biol. 215, 403–410 (1990).

21. Consortium, T. U. UniProt: a worldwide hub of protein knowledge. Nucleic Acids Res. 47, D506–D515 (2018).

22. Yang, J. et al. Care: a benchmark suite for the classification and retrieval of enzymes. Adv. Neural Inf. Process. Syst. 37, 3094–3121 (2024).

23. Cover, T. & Hart, P. Nearest neighbor pattern classification. IEEE Trans. Inf. Theory 13, 21–27 (1967).

24. Kuhner, M. K. & Felsenstein, J. A simulation comparison of phylogeny algorithms under equal and unequal evolutionary rates. Mol. Biol. Evol. 11, 459–468 (1994).

25. Wang, J. et al. Milvus: A purpose-built vector data management system. in Proceedings of the 2021 international conference on management of data 2614–2627 (2021).

26. Torgerson, W. S. Multidimensional Scaling: I. Theory and Method. Psychometrika 17, 401–419 (1952).

27. Maaten, L. van der & Hinton, G. Visualizing Data using t-SNE. J. Mach. Learn. Res. 9, 2579–2605 (2008).

28. McInnes, L., Healy, J., Saul, N. & Großberger, L. UMAP: Uniform Manifold Approximation and Projection. J. Open Source Softw. 3, 861 (2018).

29. Sokal, R. R., Michener, C. D., & others. A statistical method for evaluating systematic relationships. (1958).

30. Kuhner, M. K. & Felsenstein, J. A simulation comparison of phylogeny algorithms under equal and unequal evolutionary rates. Mol. Biol. Evol. 11, 459–468 (1994).

