## Supplementary material for "ProtSEC: Ultrafast Protein Sequence Embedding in Complex Space Using Fast Fourier Transform": Table S1, Table S2, Figure S1, Figure S2, Figure S3

### **Supplementary information**

**Table S1:** Characteristics of the protein sequence datasets used.

| **Dataset** | **num_seqs** | **sum_len** | **min_len** | **avg_len** | **max_len** |
| --- | --- | --- | --- | --- | --- |
| 5K | 5,000 | 1,829,271 | 11 | 365.9 | 5,098 |
| 10K | 10,000 | 3,670,870 | 10 | 367.1 | 6,298 |
| 20K | 20,000 | 7,420,286 | 10 | 371 | 9,904 |
| 40K | 40,000 | 14,884,713 | 5 | 372.1 | 12,345 |

**Table S2**: Hugging Face repository URLs for pre-trained protein language models

| **PLM** | **URL** |
| --- | --- |
| prot_bert | https://huggingface.co/Rostlab/prot_bert |
| esm2_35M | https://huggingface.co/facebook/esm2_t12_35M_UR50D |
| esm2_3B | https://huggingface.co/facebook/esm2_t36_3B_UR50D |
| prot_t5 | https://huggingface.co/Rostlab/prot_t5_xl_uniref50 |

**Figure S1:** Accuracy of ProtSEC for protein similarity searches using different dimension reduction (MDS, t-SNE and UMAP) and distance (ASMP, SMS, SNN) parameters. 5K dataset was used as a query.

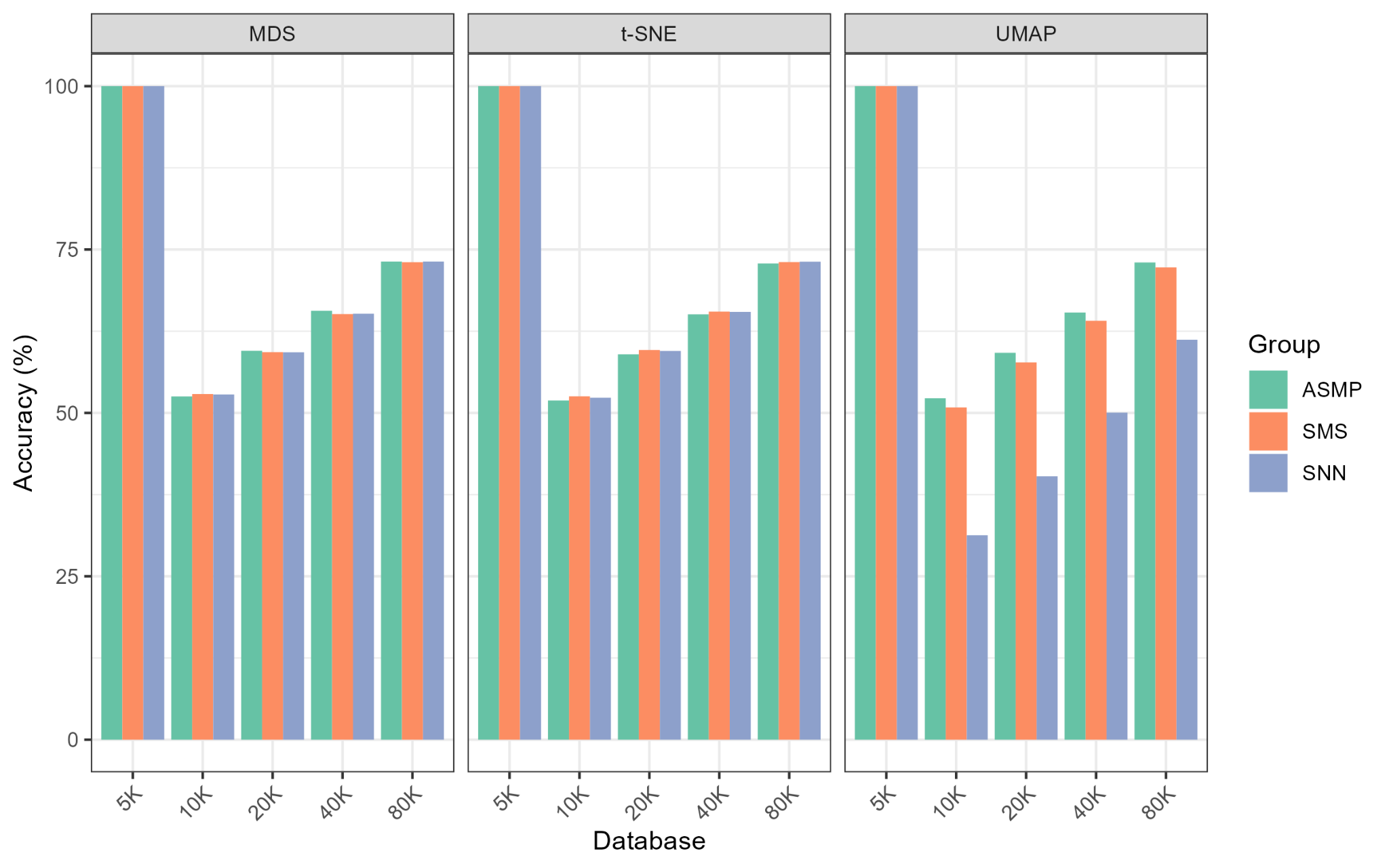

**Figure S2:** Protein sequence similarity search pipeline using PLM-based embedding

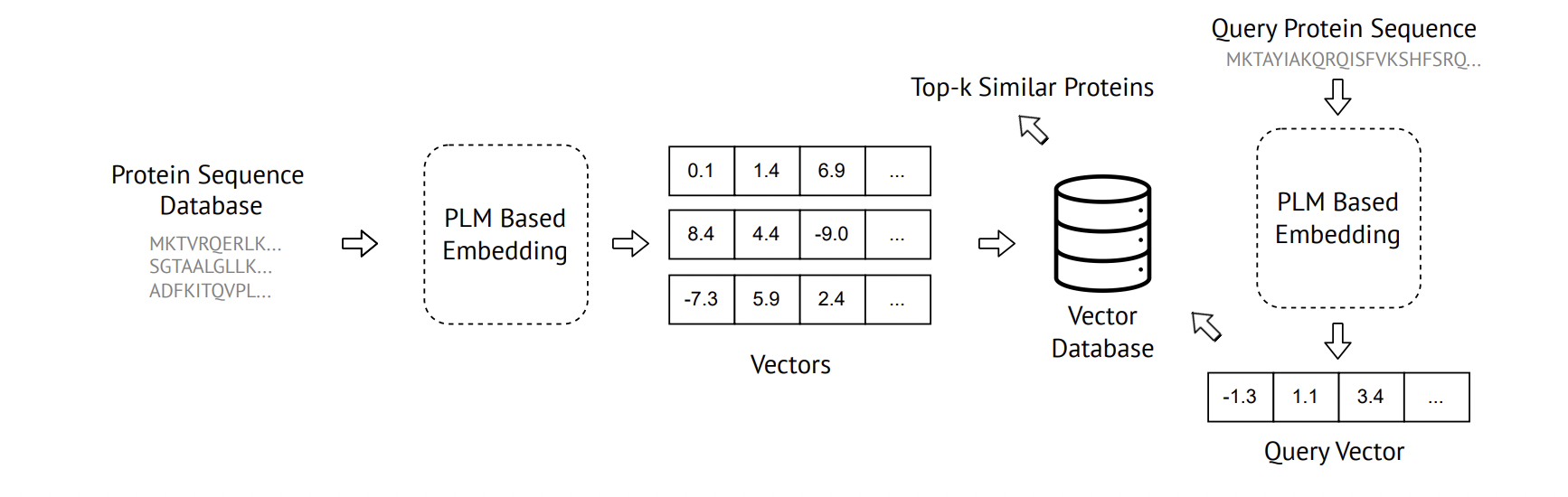

**Figure S3: Phylogenetic reconstruction and clustering performance of ProtSEC**

**A)** Phylogenetic trees of the 10-BetaSet dataset reconstructed using ClustalW (multiple sequence alignment), ProtSEC, and PLM-based embeddings (esm2_3B, esm2_35M, prot_t5, prot_bert). For BSD, trees were normalized by scaling all branch lengths proportionally.

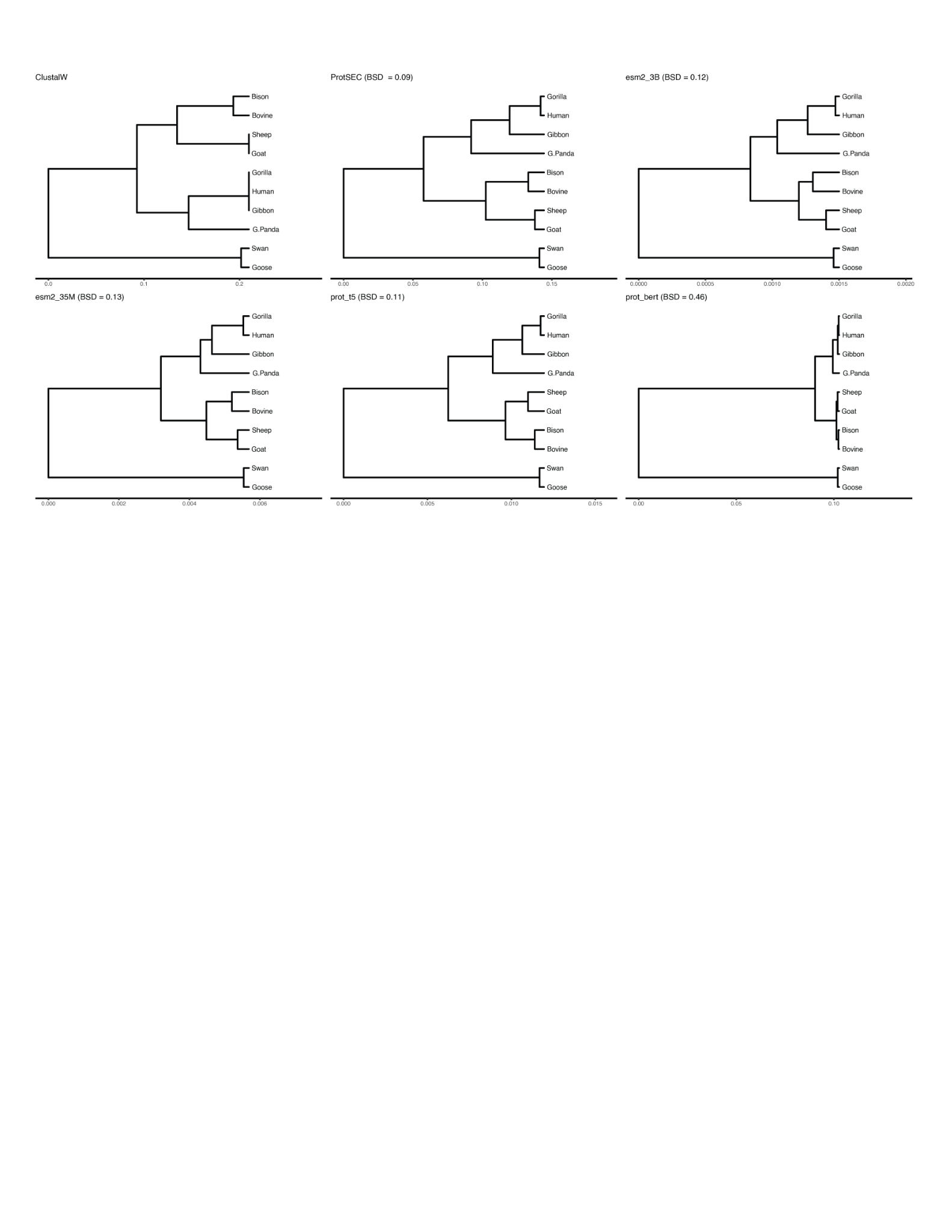
